# New modular assays for the quantitative study of skylight navigation in flying flies

**DOI:** 10.1101/527945

**Authors:** Thomas F. Mathejczyk, Mathias F. Wernet

## Abstract

The quantitative study of behavioral responses provides crucial information about how neural circuits process visual information, thereby revealing the computations responsible for shaping the animal’s perception of the outside world. Over the last decade, insects have served as particularly powerful model systems, either when walking on air suspended balls (spherical treadmill), or when flying while glued to a needle (virtual flight arena). The use of virtual flight arenas is complicated by the fact that an effective experimental setup needs to combine a rather complex set of custom-built mechanical, electronic, and software components. Assembling such an apparatus amounts to a major challenge when working in an environment without the support of a machine shop. Here we present detailed instructions for the assembly of virtual flight arenas optimized for *Drosophila* skylight navigation, which can easily be modified towards other uses. This system consists entirely of off-the-shelf parts and 3D-printed components, combining a modular flight arena designed to reduce visual artifacts, swappable high-power LED light sources, polarization filters on a computer-controlled rotating filter wheel, all placed within a temperature and humidity controlled environment. Taken together, these findings demonstrate the usefulness of these assays for the study of skylight navigation in flies.

## Introduction

The quantitative study of visual behaviors using single flies suspended in a virtual flight arena has long served as a powerful method for probing photoreceptor function, behavioral mechanisms, as well as the computational role of underlying circuit elements^1–3^. First established using larger insects, these assays have become particular useful in the dissection of visual circuitry in *Drosophila melanogaster*, by taking advantage of its unique molecular genetic toolkit^1^. Since early adaptations to *Drosophila*^*4, 5*^, virtual flight arenas have been used to quantify behavioral responses to a multitude of visual stimuli, ranging (for example) from moving edges, different colors, celestial bodies, learned shapes, to more complex visual scenes^6–12^. Visual responses of flying *Drosophila melanogaster* to linearly polarized light emanating from above (thereby simulating the celestial polarization pattern) using a flight simulator have also been demonstrated^13–15^. In these experiments, flies are presented a linearly polarized stimulus which, in many cases can be generated using commercially available polarization filters^16^. Fairly little is known about the navigational capabilities of free-living fruit flies, yet catch-and-release experiments suggest that several *Drosophila* species are able to keep straight headings over extended periods of time, while flying in desert environments which few visual landmarks (reviewed in^17^). Skylight navigation experiments using virtual flight arenas therefore serve as an attractive platform for the quantitative study of the navigation skills of both wild type insects, as well as of specimens harboring well-defined circuit perturbations.

So far, the assembly of a virtual flight arena for studying skylight navigation in flying flies required the combination of very specific skill sets ranging from the construction and assembly of both mechanical and electrical parts, as well as programming of the codes necessary for the motorized rotation of the polarization filter. In addition, tight control over ambient temperature and humidity are crucial^18^, both for reproducible wild type behavior, as well as for certain genetic perturbations. Assembly of such a complicated apparatus is a major challenge when working at a research institution without a dedicated work shop, a problem that becomes more immanent with many University departments discontinuing such services. Alternatively, the purchase of custom-made off-the-shelf options remains rather costly and cannot easily be fit into a realistic research budget. The recent development of affordable 3D filament printers combining both fast and precise printing performance with a choice of cheap and swappable printing materials for different applications now offers an attractive solution for generating new and affordable, quantitative assays for studying anatomy, behavior and physiology of different model organisms^19, 20^. Furthermore, an interactive community of researchers producing and sharing both 3D printing instructions as well as software codes for related robotic applications has now made it possible that such assays can now be introduced in a large scale in schools and Universities, both in industrial and developing countries.

Here we present detailed instructions for the assembly of temperature- and humidity-controlled virtual flight arenas optimized for, but not limited to studying skylight navigation in *Drosophila*. These instructions include: (1) a detailed list of items that can be ordered off-the-shelf (experimental enclosure, cameras, polarization filter, LED light sources, magnets, needles, heater, humidifier, etc.); (2) a detailed list of commercially available electronics/robotics components (servo motors, amplifiers, power adaptors, cables); (3) custom-made open-source code for the execution of different motor commands enabling defined rotations of the polarization filter; (4) custom-made open-source instructions for 3D-printing of all parts necessary to assemble a functioning virtual flight arena. Taken together, we estimate the cost for assembling a temperature- and humidity-controlled setup containing 2 flight simulators (in their most complete version) around 8000 EUR, assuming an existing access to a 3D filament printer. However, costs can be greatly reduced depending on experimental requirements, due to the modular concept of the setup. In order to demonstrate the usefulness of this experimental setup, we performed a series of experiments investigating the navigational decisions of individual wild type flies suspended under a rotating polarization filter. In agreement with previous studies ^14^ we find that flies choose a heading with respect to the incident e-vector field, and try to maintain this heading when the e-vector is abruptly changed. When the incident light is unpolarized, flies are not able to adjust their flight heading in response to rotations of the filter wheel. Flies show a strong tendency for maintaining their heading relative to the incident e-vector field during two consecutive trials, even when these were interrupted by a period of unpolarized light. These experiments underscore the usefulness of the experimental setups presented here and serve as an ‘open source’ platform for the development of new assays optimized for different visual behaviors, in flies as well as other species of flying insects.

## Results

The aim of this study was to develop an experimental apparatus for the study of skylight navigation of *Drosophila*, where single flies are glued to a steel needle and suspended in flight via the use of two magnets (magnetotether) ^21^ under a rotating polarization filter, while flying inside a backlit cylinder (in order to reduce reflection artifacts), being filmed by an infrared camera from below in order to extract the fly’s heading.

### New modular assays for studying skylight navigation in individual flying flies

Our new virtual flight arenas are assembled from a combination of commercially available parts and custom-designed components that can be 3D-printed using standard filament printers (Figure 1A,B) (see supplemental materials for a list and description of components and detailed building instructions). One individual assay is fixed on a 30cmx30cm double density optical breadboard (Thorlabs) via two compatible metal beams (Thorlabs, supplemental Figure 1A,B). To these, high-power LED light sources with collimated optics (Mightex) can be attached via a 3D printed, horizontal holder (upper holder). LED’s are held in place via magnets, to facilitate swapping light sources of different wavelengths. A robotics-grade servo motor for rotating the polarization filter (Dynamixel MX-28T, Robotis) is attached via a second horizontal holder (lower holder) which is attached to the rotatable filter wheel. The filter wheel contains a 3D-printed gear system as well as two removable cassettes into which polarization filters, diffusers and quarter wave plate retarders can be placed (described in more detail below), in order to produce unpolarized stimuli, or to simulate different degrees of polarization, if desired (Figure 1C). The fly is glued to a steel pin using UV-cured glue and held in place by a V-shaped sapphire bearing and a magnet (the top magnet) attached to a UV fused silica plate that rests in a horizontal platform which is in turn held in place by four vertical metal bars (Figure 1B,D). To extract the animal’s body axis, it is filmed using an infrared camera (Firefly MV, Point Grey) pointing upwards (Figure 1B,E), yet shielded from the fly’s view via a small box containing infrared LED’s, on which rests a round bottom plate containing a pinhole and the bottom magnet (see Figure 1B for magnification). The setup is completed by a white, backlit cylinder containing LED strips for reducing intensity artefacts caused by linearly polarized light being reflected off the cylinder walls (reviewed in ^16^). This cylinder can be moved vertically, either downwards to allow for insertion of a new tethered fly (see Figure 2A,B), or back upwards to shield the fly from unwanted external stimuli. Due to the placement of the upper magnet above the fly, its central dorsal field of view is somewhat restricted. Similarly to other existing assays, we calculated the stimulus to be visible within a concentric rim around the upper magnet spanning ~28° of the dorsal visual field (Figure 1E).

**Figure 1:**
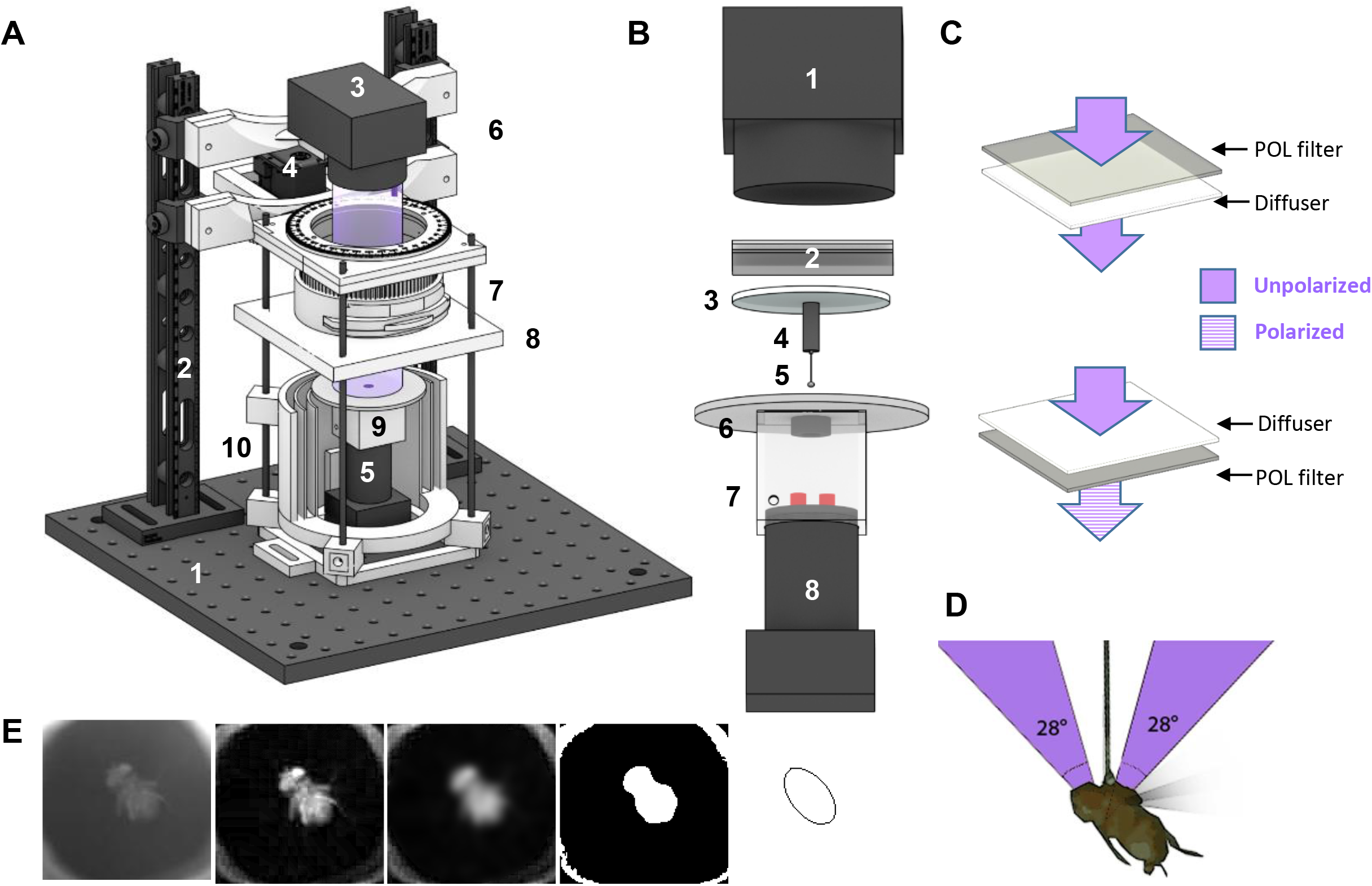
A new modular assay for studying skylight navigation in individual flying flies. **A.** Schematic drawing of a fully assembled virtual flight setup for the quantitative study of skylight navigation in flying *Drosophila*. Commercially available parts include: (1) a 30cmx30cm double density optical breadboard (Thorlabs); (2) compatible metal beams (Thorlabs); (3) swappable, magnetically attached high-power LED light sources with collimated optics (Mightex); (4) a robotics-grade servo motor for rotating the polarization filter (Dynamixel MX-28T, Robotis); (5) an infrared camera shielded from the fly’s view, filming its body axis (Firefly MV, Point Grey). Custom-designed 3D-printed parts (see supplemental materials for printing instructions) include: (6) attachment to the vertical bars; (7) filter holder including gear system (see Figures 2 and 3 for details); (8) horizontal platform holding the UV fused silica plate and top magnet for attaching the magnetotethered fly (see Figure 1B for magnification); (9) infrared illumination (enclosed LED’s) and bottom plate with ring magnet, mounted onto the infrared camera; (10) backlit cylinder for reducing linearly polarized reflection artifacts, which can be lifted up all the way to the horizontal platform, when flies are tethered. **B.** Magnification of central components; from top to bottom: (1) LED light source; (2) polarization filter and diffuser, mounted together within the rotatable filter holder (see Figures 2 and 3); (3) UV fused silica plate for attaching the magnetotether; (4) top magnet; (5) steel pin with fly attached; (6) ground plate with bottom ring magnet below; (7) encased infrared LED’s for illuminating of the fly from below; (8) infrared camera for filming the fly’s body axis. **C.** Two polarization filter / diffuser orientations can be chosen within the filter holder: Unpolarized (top), or polarized (bottom). See Figure 3B for polarimetric characterization. **D.** Drawing of a tethered fly’s field of view of the dorsal stimulus inside the apparatus. **E.** Series of images visualizing the extraction of the fly’s boy axis from camera images using ellipse fitting.

**Figure 2:**
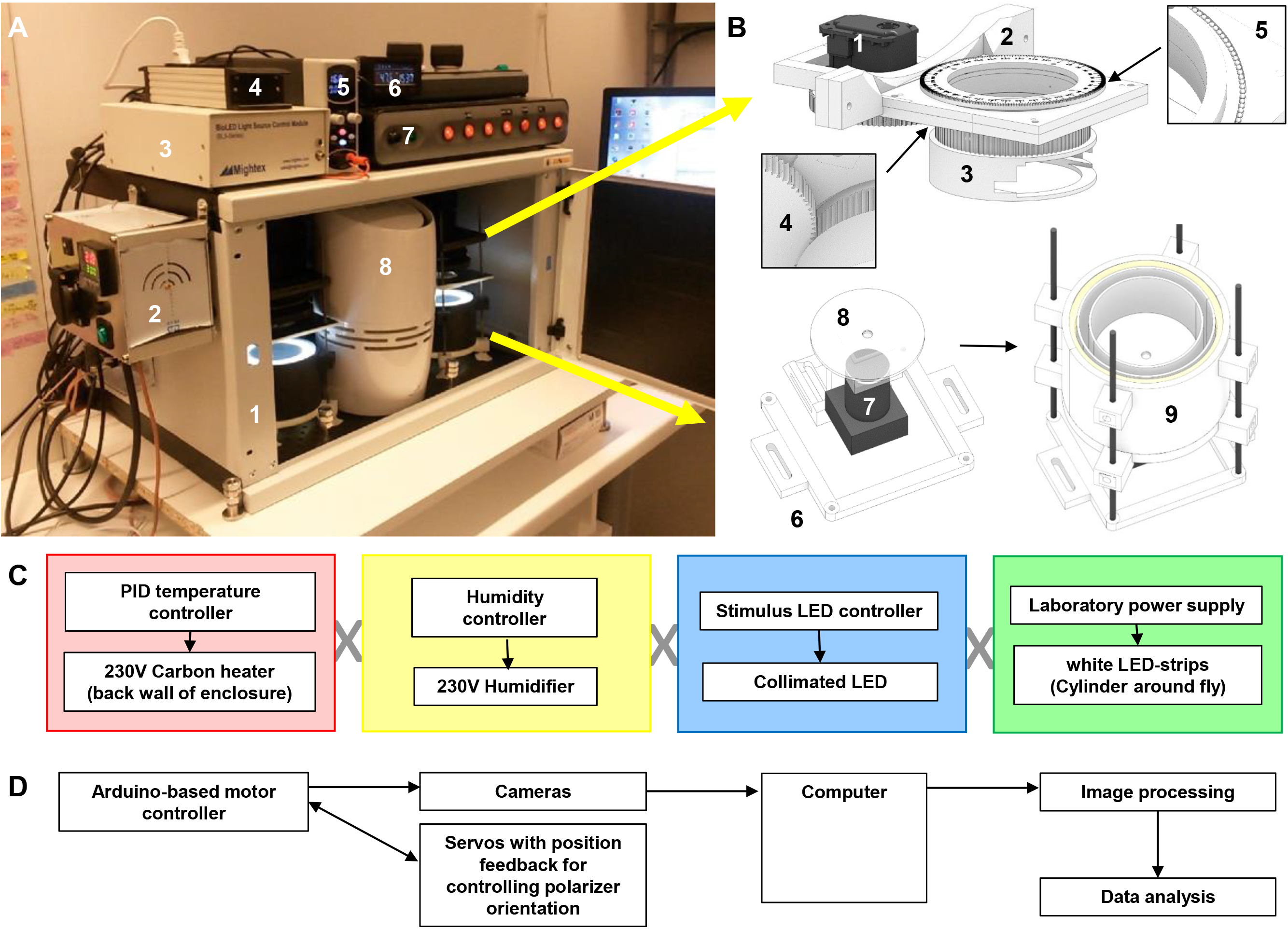
Operation of fully-assembled flight arenas. **A.** Photograph depicting two fully assembled virtual flight arenas. (1) temperature- and humidity-controlled enclosure (Monacor Rack-6W); (2) PID Temperature Controller; (3) stimulus LED controller (BLS-SA04-US, Mightex); (4) humidity controller; (5); laboratory power supply; (6) LCD display humidity controller; (7) switchable power bank; (8) humidifier (Philips HU4706/11); for a complete list of parts, see supplemental materials. **B.** Magnified view of 3D-printed parts. Top: Stimulation section. (1) servo motor; (2) attachment to vertical beams; (3) filter holder (for more details see Figure 3A); (4) gear system for rotating the filter holder; (5) ball bearing enabling smooth rotation. Bottom: modular arrangement of camera placement and optical isolation: (6) ground plate; (7) mounted infrared camera; (8) ground plate through which fly is filmed; (9) backlit cylinder containing white LED-strips for providing uniform visual environment without polarized reflections can be moved vertically around the near-infared camera / magnetotether via 4 metal guides. **C.** Summary of the four modules interacting within the apparatus (see supplemental materials). **D.** Diagram depicting information flow (For details and codes, see supplemental materials).

### Assembly of virtual flight arena from 3D-printed and off-the-shelf parts

In order to result in reproducible data, behavior experiments need to be performed in a stable environment. We therefore placed our behavioral assays within affordable enclosures (19” network rack, Monacor Rack-6W) with enough room to house up to three virtual flight arenas. However, we decided to sacrifice the central spot in favor of an evaporation humidifier (Philips HU4706/11) which can be regulated via an LCD display humidity controller (TMT-HC-210 Humidity Control II). Temperature control is achieved via a PID Temperature Controller driving a 230V carbon heating plate (#100575, Termowelt.de) with two fans mounted on the far side of the enclosure (supplemental Figure 1C). Using this simple design, the enclosure can be stably kept at high temperatures like 32°C, thereby enabling specific heat-dependent genetic manipulations^22^ (see supplementary figure 4D).

The modular design of this new virtual flight arena both simplifies assembly, and enables project-specific modifications. Upon first installation, two major parts need to be manually processed: The rotatable filter wheel (Figure 2B) can be printed as a set of 5 parts including the gear system and assembly is quick and easy. However, we recommend that small steel balls (2 mm) be inserted manually, creating a ball bearing, so that smooth turning is achieved, even at the highest speeds supported by the servo motor, thereby ensuring quick and controlled changes in filter orientation. Similarly, the white cylinder surrounding the flying fly can be 3D-printed in one piece (see supplemental Figure 2), yet an LED strip (Paulmann MaxLED 1000) needs to be inserted manually into the wall of the cylinder upon first installation.

As a whole, the virtual flight simulator consists of four independently powered control units (Figure 2C): PID temperature controller, humidity controller, stimulus LED controller, and a laboratory power supply for powering the white LED strips in the cylinder. These components can easily be coordinated via a switch board (Figure 2A). The recording setup consists of an Arduino-based motor controller (Arbotix-M) which TTL-triggers a free running recording of the two attached cameras via the recording software Streampix 7 (Norpix). Synchronously, it coordinates motor movements which rotate the pol-filter (0.088° resolution) while receiving positional feedback from the motors. The camera streams get slightly compressed using MJPEG compression and stored onto the same computer for later image processing and data analysis.

### Characterization of the visual stimulus

Important controls for testing behavioral responses to linearly polarized light include either (i) depolarizing the stimulus, while keeping its intensity constant^23, 24^, or (ii) converting the linearly polarized stimulus into circularly polarized light, which should be perceived as unpolarized, by most if not all insect photoreceptors^25, 26^. We have designed the filter holder of our experimental setup in a way that can quickly and easily accommodate for both kinds of controls, thereby making lengthy adjustments to the experimental design unnecessary. In its assembled state, the rotatable filter holder contains one of two types of removable filter cassettes (Figure 3A,C). The first type contains a shallow bed (5 x 5 cm) into which polarization filter and several sheets of diffuser can be placed. This filter cassette can easily be removed and inverted, between two experiments, thereby transforming a linearly polarized stimulus into an unpolarized one. The second filter cassette type contains a rotatable holder for quarter wave plate (QWP) retarders. Using the internal gear system, the QWP can be rotated into different fixed orientations with regard to the polarization filter, thereby producing elliptically to circularly polarized light, depending on its position. To verify the efficacy of this stimulus presentation system, we have characterized the stimuli emanating from our apparatus using a polarimetric camera^27^ (Figure 3B). By calculating the degree of linear polarization as previously described^27^, we confirm that the polarized stimuli manifest a high degree of polarization both in the UV (365 nm) as well as with a green LED (510 nm). In contrast, virtually no polarization is detected when the polarization filter / diffuser combination of the upper filter cassette is inverted.

**Figure 3:**
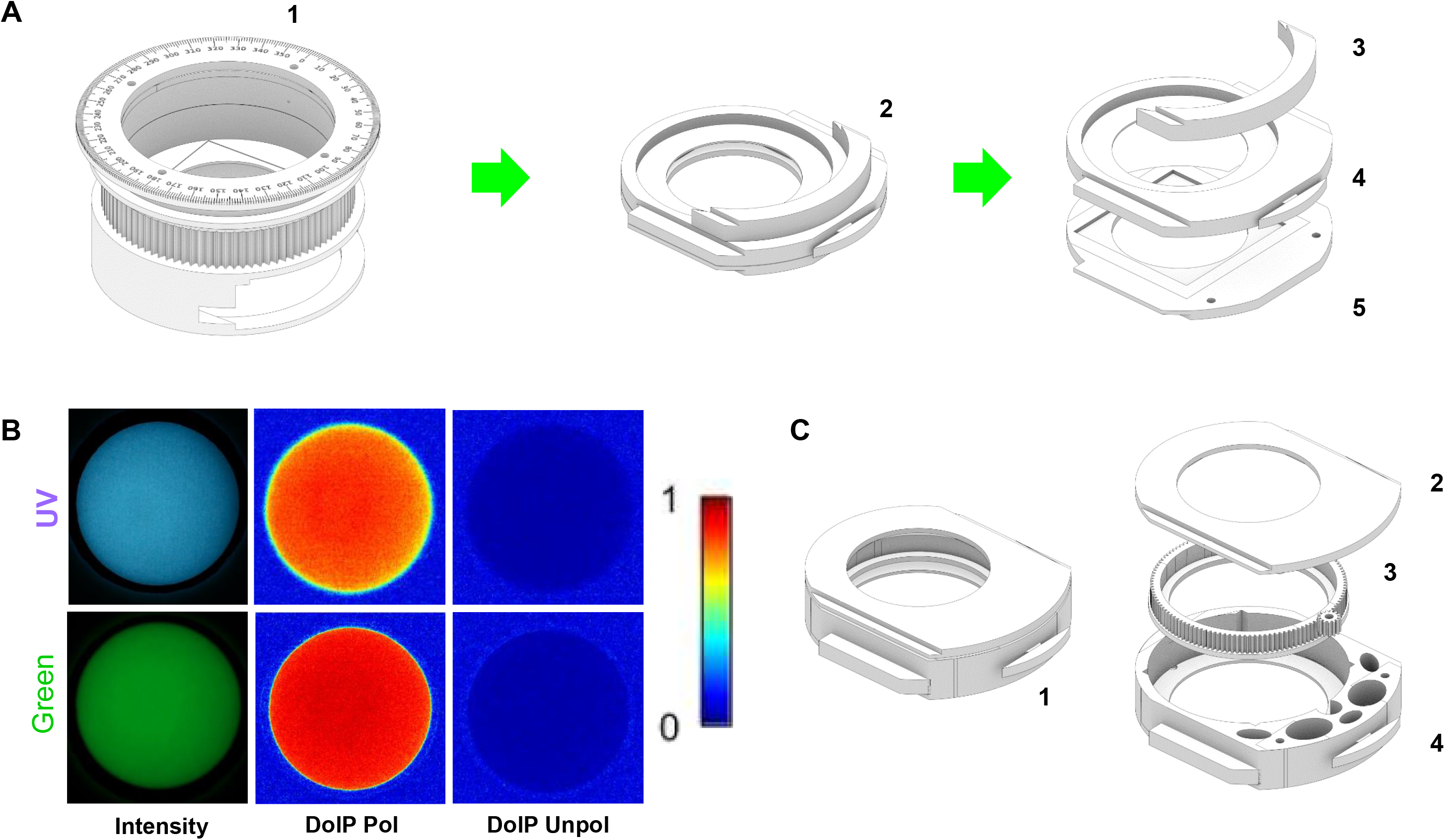
Stimulus presentation. **A.** Modular depiction of the filter holder and its individual components: (1) rotatable cassette holder; (2) Removable filter cassette; (3) optical seal; (4) round aperture; (5) filter bed (5 x 5 cm). **B.** Polarimetric characterization of stimuli as they emanate from the above apparatus. Top: UV stimulus, 365 nm (from left to right: intensity, as well as the two filter/diffuser orientations shown in Figure 1C: polarized (center) and unpolarized (right). Bottom: Same analysis for a linearly polarized green stimulus (510 nm), demonstrating the suitability of the setup for longer wavelengths. **C.** Alternative filter cassette for accommodating quarter wave plates for creating elliptically polarized light: (1) removable bottom cassette; (2) top plate of filter cassette; (3) rotatable holder for quarter wave retarder plate; (4) core of filter cassette.

### Flying *Drosophila* adjust their heading after rapid switches of the incident e-vector

As a first series of experiments we tested our virtual flight arenas by recording behavioral responses of single flying flies to rapid switches of the incident linearly polarized UV light presented dorsally (PolUV1). In a single experiment, flies were recorded for 5 minutes with the polarized UV stimulus continuously switching back-and-forth between two orthogonal filter positions (termed a and b), every 30 seconds. Upon such rapid 90° changes of the incident polarized stimulus we observed that many flies also showed the tendency to rapidly adjust their flight heading by about 90°, following the filter switch (Figure 4A). On average, a clear increase in the fly’s angular velocity was measured shortly after rapid filter switches, indicating robust behavioral responses to changes in the polarization. However, when the same fly was presented with unpolarized UV light of the same intensity as the polarized UV stimulus (UVunpol), the fly did not react to rapid filter rotation, neither by rapidly adjusting its heading, no by increased angular velocity after the switch (Figure 4B). Importantly, when the UV stimulus was repolarized by flipping the upper filter cassette (see Figure 1C and Figure 3A), the above described behavioral responses of the fly to rapid filter switches (adjustment of heading angle; increase of angular rotation) was restored (Figure 4C). When pooling data from all tested flies (N = 69) flying under the above described regime (consecutive 5 min epochs of polarized and unpolarized UV stimuli), flies showed significantly larger angular differences in flight heading between a and b periods when flying under polarized light, as compared to unpolarized trials (Figure 5A). Similarly, the increase in angular velocity shortly after rapid filter switches also differed significantly in flies flying under a linearly polarized UV stimulus, as compared to trials under unpolarized UV light of the same intensity (Figure 5B). Taken together, these findings indicate that single flies flying in our virtual flight arena indeed reacted to changes in e-vector orientation rather than intensity- or other artifacts that could possibly exist within the arena.

**Figure 4:**
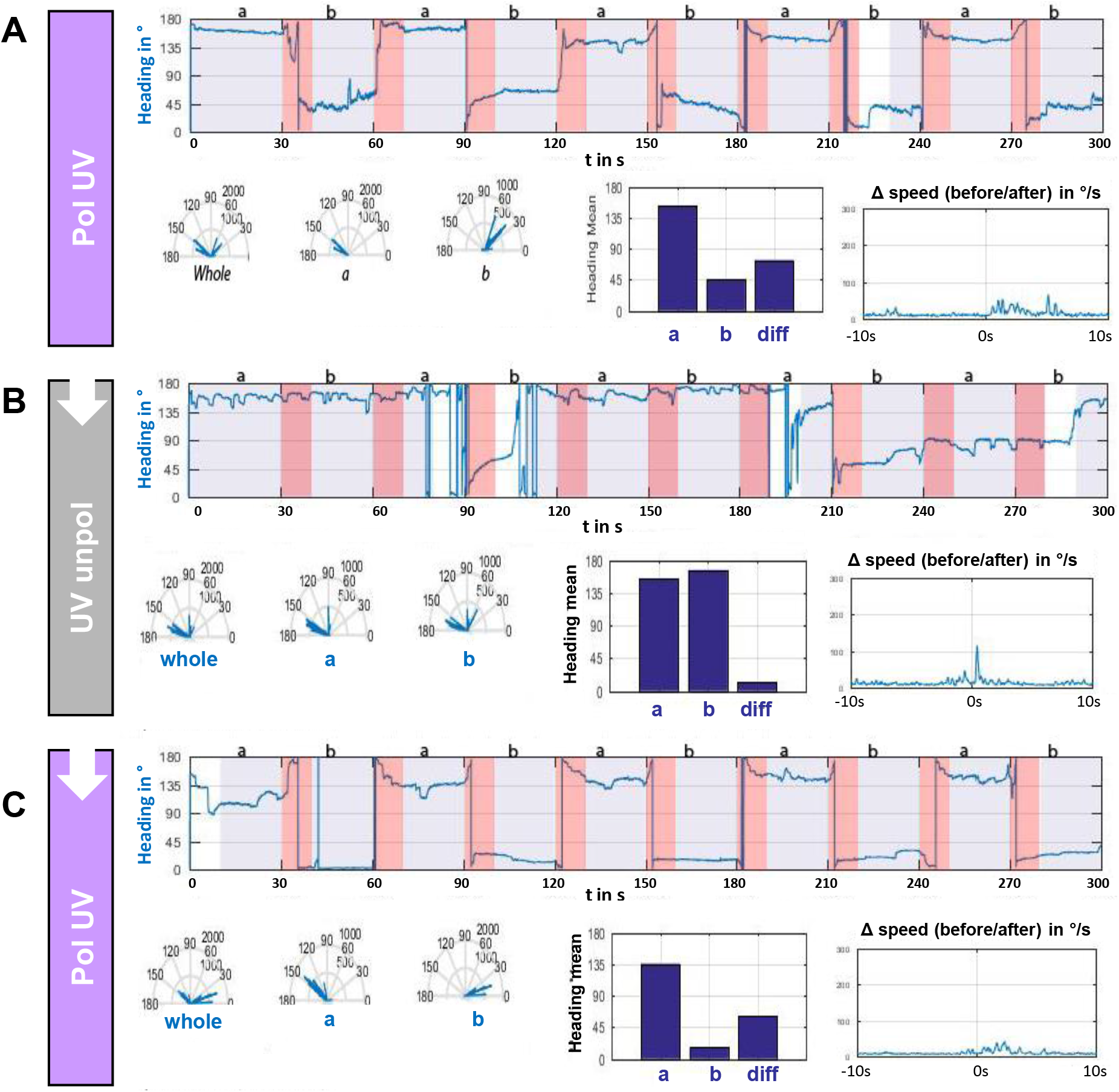
Single flying *Drosophila* react to rapid switches of the incident e-vector. **A.** Flight heading (blue line) of a single fly orienting to a linearly polarized UV stimulus (UVpol) that is rapidly changing, every 30 seconds by 90 degrees (from position ‘a’ to position ‘b’ and back). 5 consecutive series (a/b) are shown. Blue shaded areas: 10 s time windows used for analysis, where the angular speed of the fly was smaller than 0.05 deg/frame, indicating a possible response to a static e-vector. Red shaded areas: 10 s time windows after e-vector orientation switch. Bottom left: circular histogram of all heading angles (‘whole’), or separated by polarization filter orientation (‘a’ and ‘b’ periods, respectively). Bottom, center: mean heading of ‘a’ and ‘b’ periods, and difference between the two. Bottom, right: plot showing the angular speed difference (in degrees per s) 10 s before and after the rapid polarization filter switch. **B.** Same analysis for an unpolarized UV stimulus (‘UV unpol’), directly following the above experiment (same fly). **C.** Same analysis for a re-polarized UV stimulus (‘UVpol’), directly following both above experiments (same fly).

**Figure 5:**
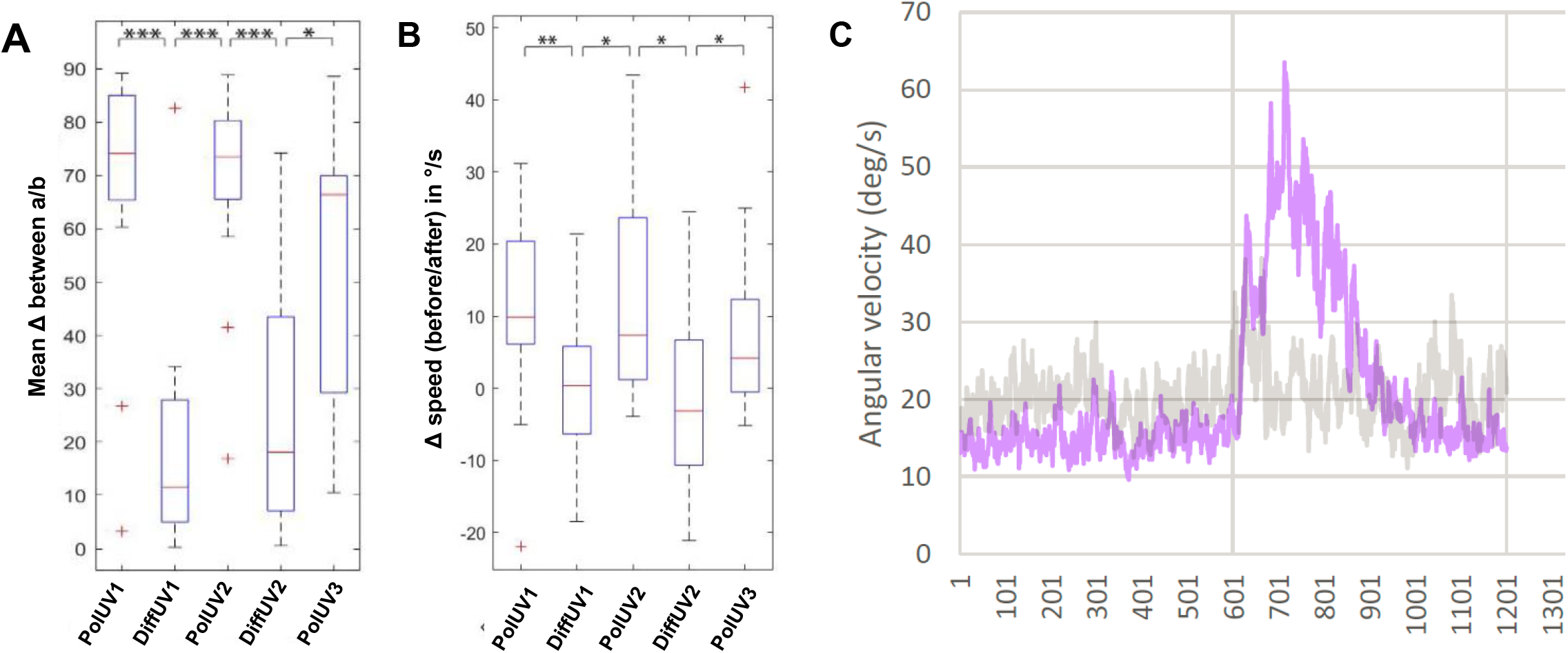
Pooled rapid switch responses from all flies tested. **A.** Mean angular difference between a and b epochs for all tested flies. The mean angular difference between a and b is significantly higher under polarized UV light compared to flies flying under unpolarized UV light. Consecutive trials alternating between polarized and unpolarized UV conditions show polarotactic responses can be elicited and prevented and that the linear polarization of light is necessary and sufficient for this behavior. *p<0.05, **p<0.01, ***p<0.001. N (left to right) = 16,16,13,13,11. **B.** Difference in angular speed between mean angular speed 10s before and 10 s after rapidly switching the e-vector for all tested flies. The difference in angular speed is significantly higher under polarized UV light compared to flies flying under unpolarized UV light. Consecutive trials alternating between polarized and unpolarized UV conditions show the necessity and sufficiency of polarized light in inducing a turning tendency shortly after rapid e-vector switching. *p<0.05, **p<0.01, ***p<0.001. N (left to right) = 16,16,13,13,11. **C.** Mean of absolute angular speed of all tested flies (grey trace: unPol, purple trace: polUV) 10s before and 10 s after rapidly switching the e-vector. Flies flying under polarized light conditions show a rapid increase in angular velocity shortly after the e-vector switch, while flies flying under unpolarized conditions do not show such changes in angular velocity upon rotating the filter wheel. *p<0.05, **p<0.01, ***p<0.001. N (left to right) = 16,16,13,13,11.

## Discussion

Navigating insects rely on the detection and integration of a combination of visual cues, like celestial bodies, intensity, gradients, and chromatic gradients^28^. In addition, the celestial pattern of linearly polarized light serves as an attractive orientation cue that many insects use^29–31^. Spontaneous behavioral responses of both walking and flying *Drosophila* to linearly polarized light (‘polarotaxis’) have been demonstrated in the past, using both population assays, as well as single fly assays^13–15, 24, 32–34^. In all these experiments, much care was given to the control and avoidance of intensity artifacts that can result in behavioral decisions that are in fact independent of the linearly polarized component of the stimulus (reviewed in^16^). However, some of the successful solutions presented in the past included components whose reproduction required considerable engineering skills and were quite costly. The virtual flight arenas presented here have been designed with the dual goal of providing relatively cheap, robust setups that can easily be assembled, while at the same time being able to provide controlled illumination, temperature and humidity conditions and minimizing intensity/reflection artifacts. For instance, reflections off the wall of any experimental chamber have been shown to produce intensity patterns that could be used by the animal as a directional cue^16^. Our setup addresses this threat by surrounding the flying fly with a white, backlit cylinder, which also serves to shield the fly from air turbulences originating from the fans within the enclosure. Furthermore, highly collimated LED light sources were chosen here in order to reduce off-axis illumination which can produce intensity confounds when hitting the polarization filter. The polarimetric characterization of the stimulus used here makes us confident that these effects have been minimized. This is supported by unpolarized UV controls, where the order of polarization filter and diffuser are inversed, which is easy and quick, using the removable filter cassette built to fit into our 3D-printed filter wheel. In addition to different combinations of polarization filters, diffusers, and band pass filters, our system can also be equipped with a quarter wave plate (QWP) / retarder that can be rotated into varying fixed orientations with respect to the polarization filter. Such an introduction of a QWP is particularly useful for testing behavioral performance while simulating different degrees of polarization^25, 26^. Although the degree of polarization in nature never reaches the high values normally used in the laboratory, data so far available only for crickets points towards the polarotactic responses to be stable across a range of degrees of polarization^25, 26, 35^.

Our compact temperature and humidity controlled enclosures provide not only stable environmental conditions crucial for most wild-type behavioral experiments, but also offer a possibiity of studying those heat-dependent genetic manipulations typically used in transgenic Drosophila, like neuronal activation via expression of TrpA1 channels, or reversible inactivation using shibire^ts 22^. Due to the modularity of the setup, enclosures of larger sizes may also be chosen, depending on experimental requirements and/or available laboratory space. Even in laboratories with access where all rooms are humidity-regulated, the humidity control intrinsic to our assay might still prove useful for humidity adjustment, for instance when performing experiments that require heating of the enclosure, since relative humidity within the enclosure will drop upon heating. Therefore, additional humidity may be required for keeping constant humidity levels during such experiments. Furthermore, when connecting a dehumidifier to the humidity controller element, it is also become possible to keep humidity levels constantly low, if desired.

Using our new virtual flight arenas, we show that individual flies choose a heading under a linearly polarized stimulus and try to maintain this heading relative to the incident angle of polarization. This finding is in good agreement with recently published studies, albeit using a rather different kind of stimulation system ^14^. In further agreement with these past studies, we also find that flies show this behavior over extended periods of time, even when interrupted by a period of unpolarized stimulation ^14, 15^. Hence, these data further support the idea that a generalist fly like *Drosophila melanogaster* is indeed capable of using skylight polarization for navigational tasks^17^. Nevertheless, our experiments also show that behavioral responses remain variable across all individuals tested. Even after tight control of food quality, rearing conditions, temperature, and humidity, the flies’ cooperation in these experiments remains unpredictable. How much this variability could depend on the fly’s motivational state or navigational decision making remains to be investigated.

We are in the process of making all the codes, templates and building instructions for the virtual flight arenas freely available for download to anyone (see materials and methods). Due to their modular design, the use of our new virtual flight assays is in no way limited to skylight navigation or the study of polarized light vision. With few simple modifications they could be modified for studying behavioral responses to moving stimuli, color patterns, or celestial bodies. Similarly, the setups can easily be modified to house a spherical treadmill, for studying the visual behavior in walking flies. Finally, application is in no way limited to just *Drosophila* or other small flies. We hope that the assays presented here can serve as a platform from which many other assays optimized for many other species could evolve from.

## Materials and Methods

### Fly rearing

An Isogenized strain of Oregon R flies^36^ was fed standard cornmeal agarose food and kept at 25°C and 60% relative humidity within a 12/12 light/dark cycle. Flies were flipped daily to ensure small population densities.

### Fly preparation

All experiments were conducted during the evening peak of the flies’ entrained activity rhythms reaching until one hour after subjective “sunset”. 3-4 days old flies were immobilized using cooling with ice and individual flies were then glued to a 1cm long 100μm diameter steel (ENTO SPHINX s.r.o., Czech Republic) pin using UV-cured glue (Bondic). The pin was attached to the fly so that when positioned vertically it held the animal at an angle pointing approximately 60° from horizontal. During the gluing procedure flies were placed onto a peltier-element set to about 4°C to keep flies immobilized. Before each experiment, flies were given at least 20 minutes to recover and were kept from flying by placing small pieces of tissue paper (Kimwipes) at their tarsi. To initiate flight for data acquisition a small air puff was given to the flies immediately from below using bellows. Flies that stopped flying more than 3 times within one trial of 5 minutes were excluded from later analysis. All experiments were conducted under 25°C and 50% relative humidity.

### Flight simulator setup

In order to study the skylight navigation in flying *Drosophila melanogaster* we modified the setup by Weir and Dickinson^14^. We created a highly modular, easy to replicate behavioral assay and in order to increase the rate of data output, placed two identical copies of this assay into one temperature- and humidity-controlled enclosure. Most functional parts of this setup were 3D printed which allows for easy replication of the setup, when having access to a 3D printer. For a complete overview of the assembly see supplemental material.

### Temperature- and humidity-controlled optical enclosures for creating constant environmental conditions

As a compact optical enclosure for our assays a 19” network rack (Monacor Rack-6W) was chosen. All removable components except the door were removed and any holes in the enclosure were sealed with aluminum tape. Two large 230mm diameter computer fans were placed in the back of the enclosure, constantly blowing at a carbon heater plate (#100575, termowelt.de). This heater plate is controlled by a PID temperature controller (RT4-121-Tr21Sd, pohltechnic.com) which was attached to the outside of the enclosure. Furthermore, an evaporation humidifier (Philips HU4706/11) was added to the enclosure (see supplemental figure 4A,C), which is controlled by an external humidity controller (TMT-HC-210, top-messtechnik.com). The sensor of the temperature controller, as well as from the humidity controller were placed close to one of the two LED cylinders. This simple and modular approach allowed us, to keep the temperature constant at ~25°C and relative humidity at ~50% during experiments.

### Fly tethering

The setup allows flies to be positioned within an axially-symmetric magnetic field created by magnets above (cylinder magnet, 4x 4mm x 5mm) and below (ring magnet, 10mm x 5mm, inner diameter 5mm) the fly, allowing the animal to freely rotate around its yaw axis. The side of the upper magnet that was facing the fly was covered with a small white matte plastic cap. The upper magnet was held in place at the center of a 50mm diameter UV fused silica Window by placing another small magnet (cylinder magnet, 5mm x 5mm) at the other side of the window. A small sapphire jewel bearing was attached to the center of this cap. This bearing served to hold the tip of the steel pin with the fly in place. The lower magnet was centrically glued to the bottom of a rough white plastic platform, hiding the magnet from the view of the fly. A 5mm hole at the center of this bottom plate allowed for delivering air puffs and also recording the fly through that hole.

### Image acquisition

For imaging the fly’s rotation and determining its heading, a camera suitable for fast near-infrared imaging (Firefly MV, Point Grey) was placed upwards so that it imaged through the hole in white bottom platform. The camera’s objective was attached to a small chamber at the bottom of the base plate. This chamber served as a spacer to increase the distance between the camera and the fly in order to match the focal range of the objective relative to the position of the fly. Furthermore, a 5mm diameter hole at the side of this chamber allowed for connecting the bellows and giving air puffs to the animal. Also, this chamber served as a housing for four infrared LEDs (880nm, 100mA max) which were placed directly onto a long-pass filter (#87C, Lee filters) centered on top of the objective lens with putty, but avoiding a small region directly at the center of the objective, allowing for the fly still to be imaged through that hole. The infrared light from the LEDs projected upwards through the hole in the bottom base plate, illuminating the fly for proper imaging while being invisible to the fly itself.

### Stimulus delivery

Above the UV fused silica window which holds the upper magnet in place, a custom designed freely rotatable Filter holder was installed. This holder allows for insertion of two tightly fitting filter cassettes. The upper cassette held a combination of a 50mm x 50mm sheet linear polarizer (OUV5050, Knight Optical, UK) and 13 layers of thin, non-fluorescent diffuser paper (80g/sqm, Max Bringman KG) and could be inserted into the rotatable cassette holder with either the polarizer or the diffuser side facing the fly. This setup allowed for presenting the fly either linearly polarized or unpolarized light dorsally, depending on which side of the filter cassette was facing the fly, providing a way to alter the degree of polarization while maintaining the light intensity. The rotatable filter cassette holder was held in place by a manually assembled ball bearing. This was done by modelling a 2.15mm gap between the filter cassette holder and its surrounding frame when constructing the 3D models. This gap was sparsely lubricated, almost completely filled with 2mm diameter steel balls and covered with a fitting top plate to keep the balls in place. The rotatable filter cassette holder contained a gear system (template created using https://geargenerator.com) which was driven by a 360° servo motor capable of sending positional feedback data (MX-28T, Dynamixel). The servo was controlled with an Arduino-compatible microcontroller (Arbotix-M, Trossen Robotics) allowing for precise rotations of the filter cassette holder and with it the angle of the e-vector of the linearly polarized light stimulus. To illuminate the fly through the polarizer/diffuser either a collimated UV (365nm, LCS-0365-13-B, Mightex) or a collimated green (530nm, LCS-0530-15-B, Mightex) LED were mounted centrally above the filter cassette holder, projecting light through the filters and to the dorsal side of the fly’s eye. The intensity of the two LEDs from the flies’ position was set approximately isoquantaly to 2×10^12^ photons/s/cm2 using a spectrometer (Flame, Ocean Optics). Since the original cooling fans of the stimulus LEDs were quite noisy and since LED intensities more than 20% were never used, much more quiet fans were installed (Sunon MB40201V3-0000-A99), in order to reduce unwanted vibrations. Hence, if higher light intensities were to be required, LED cooling power should be increased accordingly to prevent damaging the LEDs.

### Motor control of filter rotation

Data acquisition was synchronized using the Arbotix-M microcontroller which controlled motor position and triggered the infrared camera. For recording videos, we used Streampix software (Norpix). With this software it was possible to monitor the input state of one of the digital I/O connections of the camera to trigger and end a free running recording. The input state was defined by a TTL signal emanating from the Arbotix-M in order to synchronize video acquisition with motor movements. Images were acquired over 5 minutes with 60 frames per second in greyscale and 640×480 pixel resolution and stored in M-JPEG compressed AVI files. The stimulus protocol consisted of rapidly rotating the e-vector by 90°, keeping it static for 30s and then rapidly switching it back by 90° to its original position. The duration of a 90° switch was approximately 1s. This rotational pattern was repeated 10 times, resulting in a duration of recording of ~5min

### Extraction of flight heading

Tracking of the fly’s body axis angle was done offline using the open-source software Fiji ^37^. To automate this process a macro was created for Fiji which opens all relevant AVI files in a folder, separates the fly from the background, performs per-frame tracking and saves the tracked angular data of the fly’s heading to a text file. The macro involves an automated rolling ball background subtraction, a median filter to smooth out pixel noise and a thresholding command to binarize the image and separate the fly’s body from the background. Afterwards, Fiji fitted an ellipse around the fly’s body in each frame, outputting the angular data of the heading of the fly over time from 0° to 180°. Due to the directional ambiguity of the E-vector we also calculated the E-vector angle within a range from 0° to 180°. The text files containing the tracking results for each frame of a video were later analyzed in MatLab using circular statistics. In short, each track was split up into 10 s windows and only those windows where the angular speed of the fly was smaller than 0.05 deg/frame (indicating a possible response to a static e-vector) were used for analysis. Furthermore, mean headings within those periods were calculated. 10 s windows directly after switching were not used for mean heading computations, unless noted otherwise.The commented Matlab and FIJI scripts used in this study for data analysis are available for download under www.flygen.org/skylight-navigation.

### Open access

Files necessary for 3D-printing flight simulator parts, as well as codes and further assembly instructions can be downloaded under www.flygen.org/skylight-navigation.

## Acknowledgements

The authors thank Tanja Heinloth, Leigh Moss, Lena Naber and Nurelhoda Abdel Muti for experimental support, as well as all members of the Wernet and Hiesinger groups for their input. This work was supported by DFG grant WE5761/2-1 and collaborative research grant SFB958 (Teilprojekt A23), the US National Institute of Health (grant to Robin Hiesinger; division of Neurobiology), the Fachbereich Biologie, Chemie und Pharmazie der Freien Universität Berlin, the Berlin-based excellency cluster NeuroCure, as well as AFOSR grant FA9550-19-1-7005.

## Author contributions

TM and MFW planned the experiments. TM built the assay and performed all experiments. TM and MFW designed the figures. MFW wrote the manuscript, MFW and TM finalized the manuscript.

