## Supplemental Material for "New modular assays for the quantitative study of skylight navigation in flying flies"

#### Supplemental Figure Legends

Supplemental Figure S1: Summary of 3D-printed and off-the-shelf structural parts

**A.** Off-the-shelf structural components: (1) Dovetail optical rail (RLA300/M, thorlabs.com), (2) Dovetail rail spacers (SC1/M, thorlabs.com), (3) Dovetail rail carrier (RC1, thorlabs.com), (4) Mounting base (BA2/M, thorlabs.com), (5) Aluminum breadboard (MB3030D/M, thorlabs.com). **B.** Depiction of all custom-designed parts for which 3D-printable STL files are provided for download at www.flygen.org/skylight-navigation. Files labeled with an asterisk are required to be printed with white material to reduce polarization-induced intensity artefacts. We recommend to print all other parts with black material to reduce light scattering throughout the material.

Supplemental Figure S2: Assembly Instructions

Parts for self-printing are labeled with “p” and off-the-shelf items for purchase are labelled with “b”. Item numbers refer to the numbers depicted in supplemental figures 1B (p) and 3 (b), respectively. Also, filenames of the 3D files which can be downloaded at www.flygen.org/skylight-navigation follow the same numeric scheme for ease of use. **A.** Rail system. (Place breadboard into enclosure and fix rails with screws onto breadboard) **B.** Camera. (Make sure to place the NIR-LED cables so that they will not interfere with the movable cylinder depicted in (C). **C.** Movable cylinder. (After placement of white-LED strips the two cylinder parts may be glued together for better stability, when cylinder is moved upwards during experiments, it might be supported from below using a combination of knurled screws and glued hex nuts (b14, b15) as adjustable stands.) **D.** Top magnets and needle bearing. (Make sure to assemble parts as centered and lined up as possible to ensure proper placement of tethered flies within the magnetic field.) **E.** Rotatable filter wheel. (Add a little bit of grease when inserting the 2mm steel balls into the bearing. After the steel balls are placed, the top cover can also be glued to the bottom part to prevent balls from falling out. Screw the gear wheel to the shaft of the motor and ensure proper grip of the gears before fixing the motor. Connect the motor to the Arbotix-M controller.) **F.** Stimulus LED. (Screw the LED to the holder and screw the holder onto the rail carriers. Optionally, instead of using screws, magnets can be attached to LED and holder to allow for quick changes of stimulus LED and therefore wavelength.). **G.** Filter cassettes. (Insert polarizer/diffuser/quarter waveplate into filter cassettes. Top and bottom parts of the cassettes might be fixed together using tape or glue to prevent the cassette from opening.)

Supplemental Figure S3: List of off-the-shelf components

List of all flight simulator parts that are commercially available (for instructions, see materials and methods). Complete Excel list is currently being made available through www.flygen.org/skylight-navigation.

Supplemental Figure S4: More details on the experimental setup

**A.** Photo zooming in onto the two virtual flight arenas placed within the temperature- and humidity-controlled enclosure. Left: Green stimulus, right: UV stimulus. **B.** Characterization of LED light sources used (Mightex), see materials and methods. **C.** Photo of two experimental setups without humidifier, revealing the far side of the enclosure: (1) heating plate; (2) fans. **D.** Graph depicting the dynamics of temperature control in the closed setup.

### Supplemental Figure S1

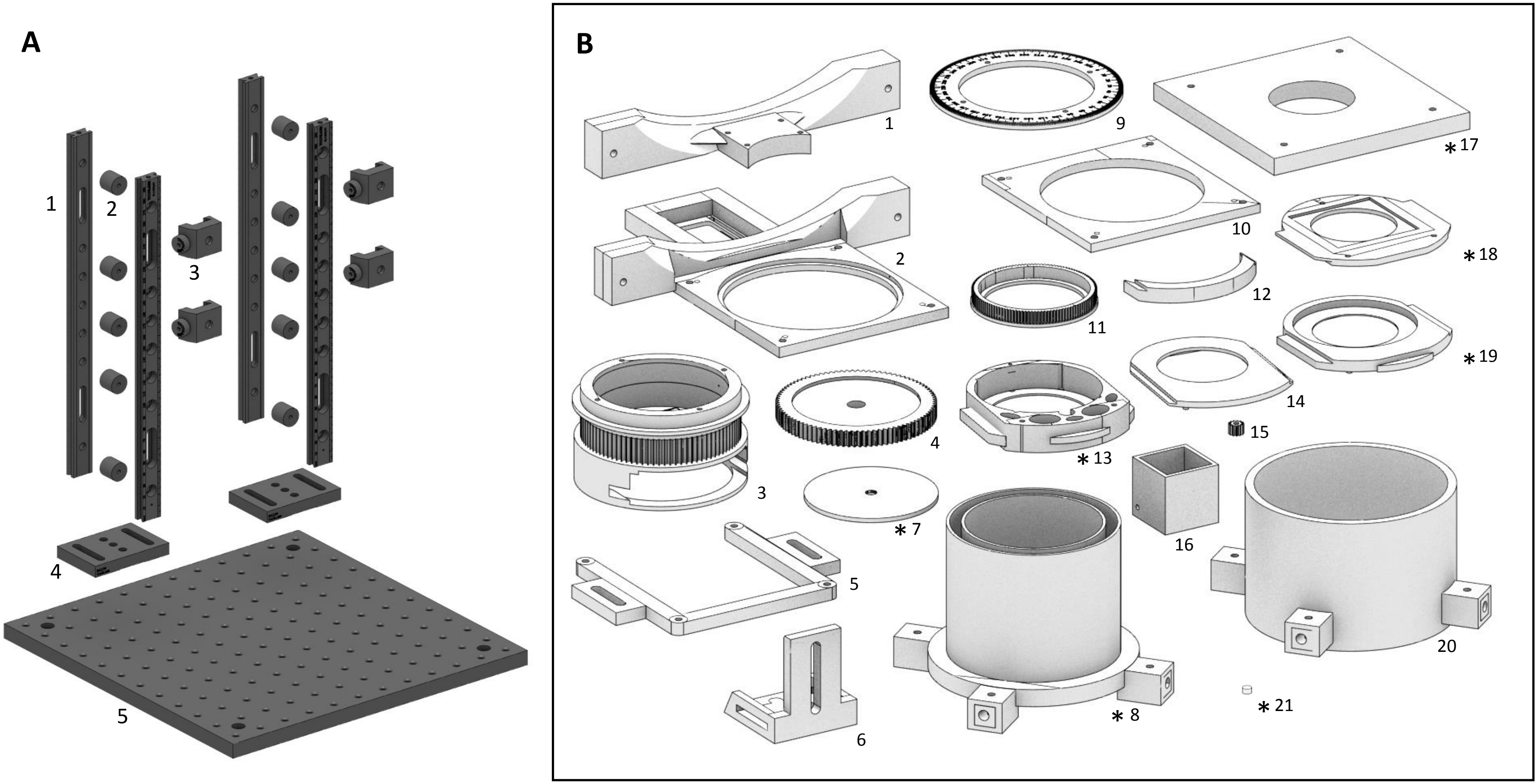

### Supplemental Figure S2

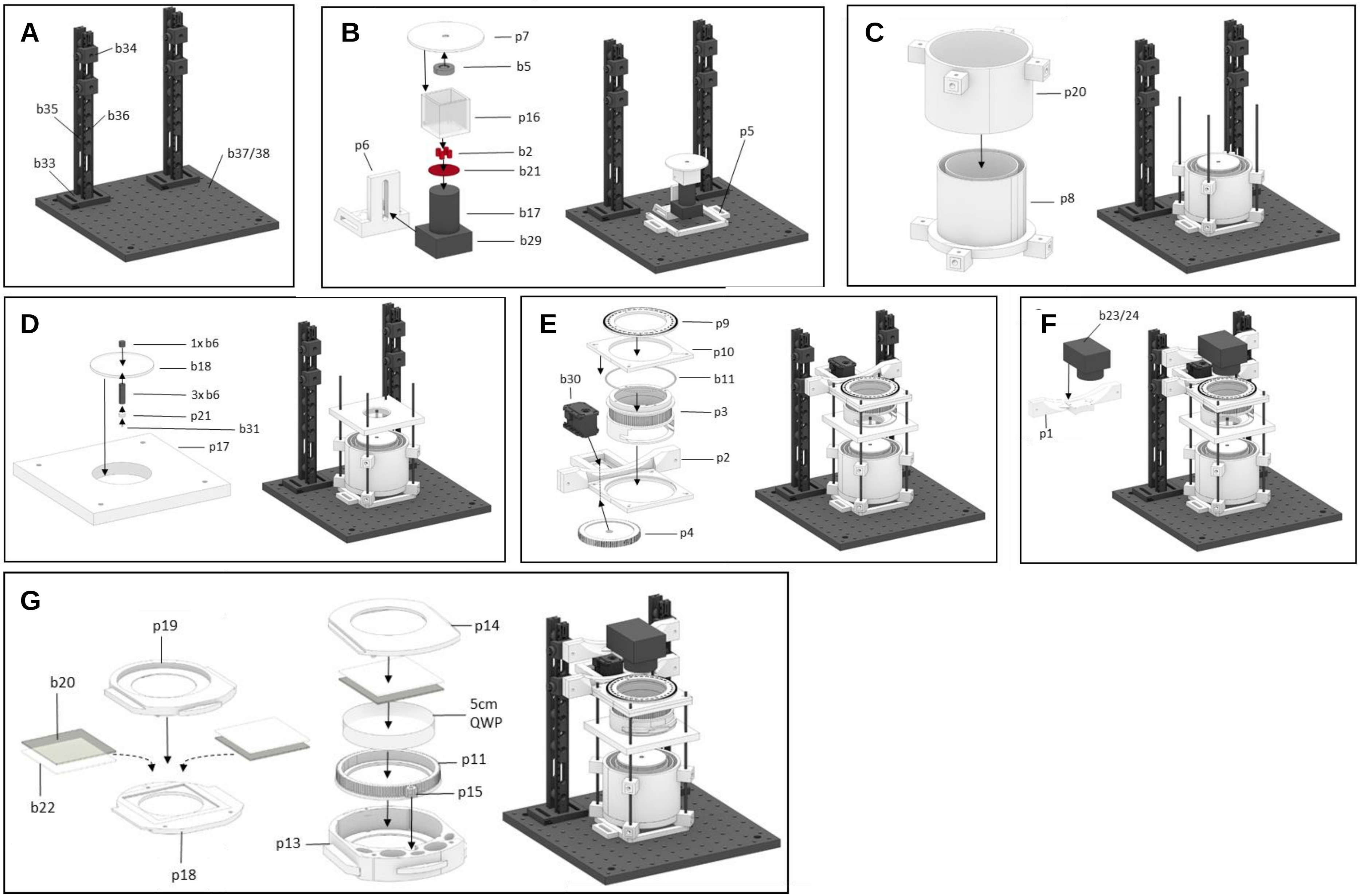

### Supplemental Figure S3

| Part # | Amount | Article | Seller/Manufacturer | Part ID |
|---|---|---|---|---|
| 1 | 1 | UV-cured glue | shop.bondic.de | - |
| 2 | 8 | Near-infrared LEDs 880nm, 100mA max | conrad.de | 153831 - 62 |
| 3 | 1 | Humidifier Philips HU4706/11 | conrad.de | 1378450 - 62 |
| 4 | 2 | Laboratory power supply VOLTCRAFT LSP-1165 | conrad.de | 1337856 - 62 |
| 5 | 2 | Ring magnet TRU COMPONENTS N35 | conrad.de | 1571358 - 62 |
| 6 | 8 | Cylinder magnet StandexMeder Electronics | conrad.de | 1205898 - 62 |
| 7 | 1 | Switchable power bar Boos RC7 | conrad.de | 971821-62 |
| 8 | 1 | Enclosure Monacor Rack-6W | conrad.de | 1332465 - 62 |
| 9 | 1 | Power strip 6x Brennenstuhl | conrad.de | 1434107 - 62 |
| 10 | 2 | 230mm fan | conrad.de | 1093404 - 62 |
| 11 | 40 | Packs of 2mm diameter steel balls | conrad.de | 220318 - 62 |
| 12 | 2 | 300cm Paulmann MaxLED 1000 | conrad.de | 1406643 - 62 |
| 13 | 4 | Threaded rods M4 | conrad.de | 1066496 - 62 |
| 14 | 1 | Pack of knurled screws m6 | conrad.de | 112086 - 62 |
| 15 | 1 | Pack of hex nuts M6 | conrad.de | 1065032 - 62 |
| 16 | 1 | Pack of hex nuts M4 | conrad.de | 1060795 - 62 |
| 17 | 2 | 8mm UC Series objective | edmundoptics.de | 33302 |
| 18 | 2 | Anti-reflex coated window UV-VIS CTD TS | edmundoptics.de | 84479 |
| 19 | 1 | Pack of 1cm long 100µm diameter steel pins | entosphinx.cz | 3.1 |
| 20 | 2 | 50mm x 50mm sheet linear polarizer | knightoptical.com | OUV5050 |
| 21 | 1 | Near-infrared filter | leefilters.com | 87C |
| 22 | 1 | Pack of non-fluorescent diffuser paper 80g/sqm | folia.de | - |
| 23 | 2 | Collimated UV LED 365nm | mightexsystems.com | LCS-0365-13-B |
| 24 | 2 | Collimated green LED 530nm | mightexsystems.com | LCS-0530-15-B |
| 25 | 1 | Stimulus LED controller | mightexsystems.com | BLS-SA04-US |
| 26 | 1 | Streampix 7 Multi camera software | norpix.com | - |
| 27 | 1 | PID temperature controller | pohltechnic.com | RT4-121-Tr21Sd |
| 28 | 1 | Temperature sensor | pohltechnic.com | PT100-13-50x6perf2S |
| 29 | 2 | Firefly MV camera | ptgrey.com | FMVU-03MTM-CS |
| 30 | 2 | 360° servo motor Dynamixel MX-28T | robotis.us | 902-0067-000 |
| 31 | 2 | V-shaped saphire bearing 1.2mm diam. | situs-tec.de | - |
| 32 | 1 | Carbon heater 230V | termowelt.de | 100575 |
| 33 | 4 | Mounting base, 50 mm x 75 mm x 10 mm | thorlabs.com | BA2/M |
| 34 | 8 | Dovetail rail carrier, 1.00" x 1.00" | thorlabs.com | RC1 |
| 35 | 4 | Dovetail rail spacers, 5 pack, M4 taps | thorlabs.com | SC1/M |
| 36 | 8 | Dovetail optical rail, 300 mm, metric | thorlabs.com | RLA300/M |
| 37 | 1 | Aluminum breadboard, 300 mm x 300 mm M6 | thorlabs.com | MB3030D/M |
| 38 | 1 | Aluminum breadboard, 250 mm x 300 mm M6 | thorlabs.com | MB2530/M |
| 39 | 1 | Humidity controller TXG | top-messtechnik.com | TMT-HC-210 |
| 40 | 1 | Microcontroller Arbotix-M | trossenrobotics.com | IL-ARBOTIXM |

This list includes parts for building two flight assays within one enclosure. Cables, tools and screws are not included.

### Supplemental Figure S4

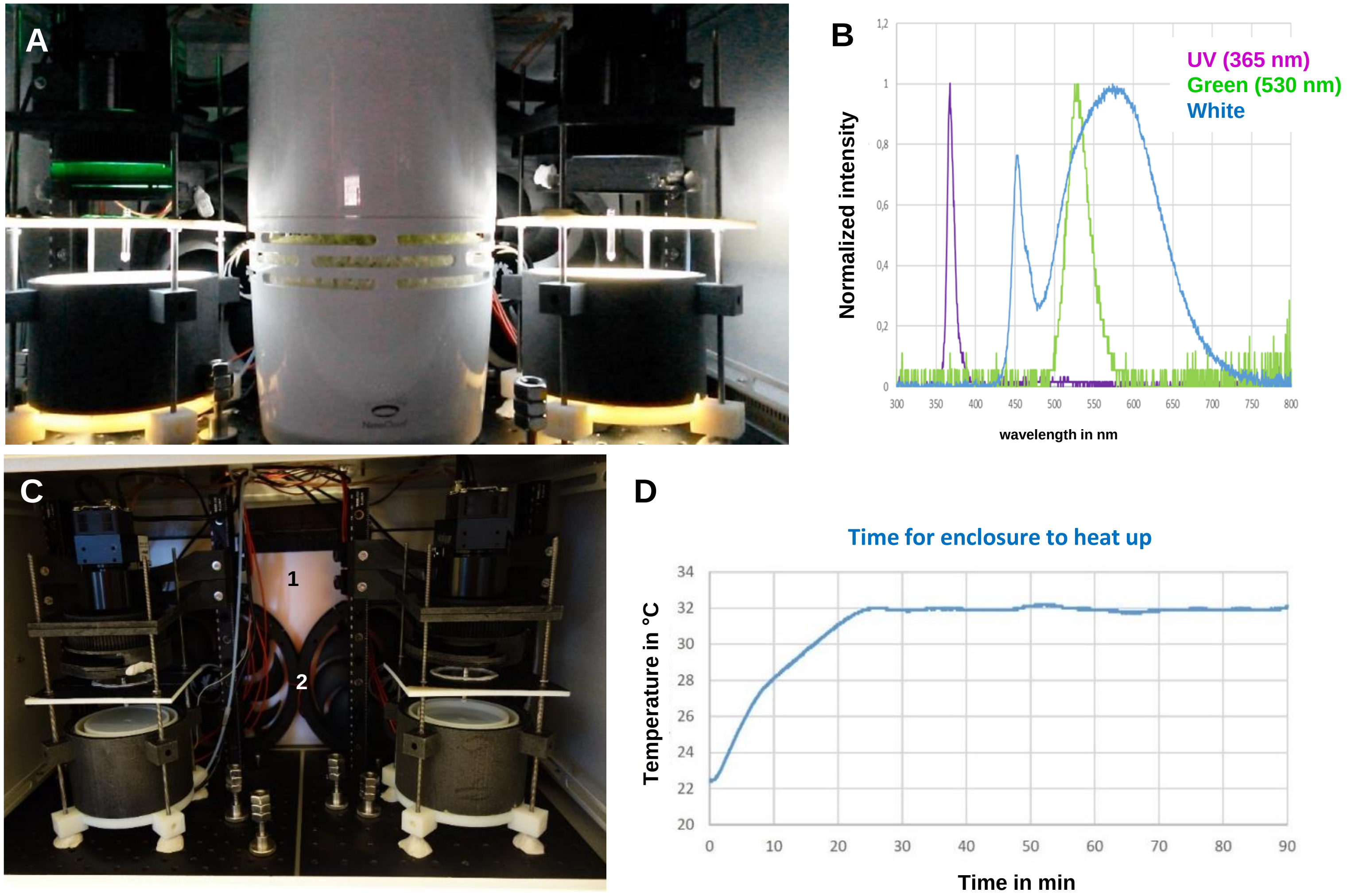
